# GoldPolish-Target: Targeted long-read genome assembly polishing

**DOI:** 10.1101/2024.09.27.615516

**Authors:** Emily Zhang, Lauren Coombe, Johnathan Wong, René L Warren, Inanç Birol

**Affiliations:** Canada’s Michael Smith Genome Sciences Centre, BC Cancer, Vancouver, BC V5Z 4S6, Canada

## Abstract

**Background:** Advanced long-read sequencing technologies, such as those from Oxford Nanopore Technologies and Pacific Biosciences, are finding a wide use in *de novo* genome sequencing projects. However, long reads typically have higher error rates relative to short reads. If left unaddressed, subsequent genome assemblies may exhibit high base error rates that compromise the reliability of downstream analysis. Several specialized error correction tools for genome assemblies have since emerged, employing a range of algorithms and strategies to improve base quality. However, despite these efforts, many genome assembly workflows still produce regions with elevated error rates, such as gaps filled with unpolished or ambiguous bases. To address this, we introduce GoldPolish-Target, a modular targeted sequence polishing pipeline. Coupled with GoldPolish, a linear-time genome assembly algorithm, GoldPolish-Target isolates and polishes user-specified assembly loci, offering a resource-efficient means for polishing targeted regions of draft genomes.

**Results:** Experiments using *Drosophila melanogaster* and *Homo sapiens* datasets demonstrate that GoldPolish-Target can reduce insertion/deletion (indel) and mismatch errors by up to 49.2% and 53.4% respectively, achieving base accuracy values upwards of 99.9% (Phred score Q>30). This polishing accuracy is comparable to the current state-of-the-art, Medaka, while exhibiting up to 36-fold shorter run times and consuming 94% less memory, on average.

**Conclusion:** GoldPolish-Target, in contrast to most other polishing tools, offers the ability to target specific regions of a genome assembly for polishing, providing a computationally light-weight and highly scalable solution for base error correction.

**Availability:** https://github.com/bcgsc/goldpolish

## Background

The genomics revolution has led to a surge in the development of bioinformatic technologies aimed at generating high-quality *de novo* genome assemblies that are crucial for extracting biological insights from sequencing data (1). Long-read sequencing, notably using Oxford Nanopore Technologies Plc. (ONT, Oxford, UK) and Pacific Biosciences Inc. (PacBio, Menlo Park, USA) instruments, has gained prominence due to its ability to capture long-range genomic information within single reads, enabling the resolution of complex structural variants and regions with high homology (2). Unlike Illumina, Inc. short-read sequencing that can generate reads up to 600 bp (3), ONT produces reads with average lengths of 10-100 kbp (4) and PacBio HiFi sequencing produces reads lengths averaging 10-25 kbp long (5). While long-read technology facilitates the generation of highly contiguous genome assemblies, its associated error rates tend to be higher (1-15%) (4,6) compared to Illumina short-read sequencing (∼0.1-1%) (7). These erroneous reads can negatively impact the base quality of subsequent genome assemblies, hindering accurate downstream analysis, such as the identification of true variants (8) or comparative genomic studies (1).

In response, specialized error correction tools, such as Racon (9), Medaka (10), and GoldPolish (11), have emerged, using long reads to identify and correct base errors in a genome assembly (12). While these tools ultimately share the same goal, the approaches of these polishing algorithms vary widely (13). Racon uses information from mapped reads to construct a partial-order alignment graph to generate a final consensus sequence (9). Medaka generates a consensus sequence by applying neural networks to sequencing reads that are aligned to a draft assembly (10). GoldPolish uses a long-read adaptation of the ntEdit+Sealer protocol (14) implemented in the GoldRush genome assembly pipeline (11). The method described in this work builds on the GoldPolish algorithm.

Many assembly algorithms leave regions of the genome with highly concentrated error rates, such as the terminal overlapping regions of fragmented contigs, in the final assembly output (15). These regions can negatively impact the reliability of downstream analyses of genome assemblies and should be addressed prior to analysis. For example, in a study conducted in 2019, human ONT long reads assembled with Canu (16), a single molecule sequence assembler, exhibited regions with higher error rates that substantially impacted the base quality of protein-coding genes, ultimately hindering protein prediction (17). Similarly, the GoldRush genome assembly pipeline can leave regions with relatively higher error rates. Specifically, during the scaffolding stage of GoldRush, instead of introducing ambiguous bases, gaps in the genome assembly are filled with unpolished (raw) bases from representative long reads and are subsequently soft-masked (i.e. lower-cased) (18). These filled-in gaps are never polished within the GoldRush workflow. To address these potential challenges that many assembly algorithms face, we developed GoldPolish-Target (GP-Target), a long-read targeted polishing pipeline based on GoldPolish.

GP-Target enables the correction of specific genomic regions without having to polish entire assemblies. With experiments using *Drosophila melanogaster* (fruit fly) and *Homo sapiens* (human) datasets, we demonstrate that GoldPolish-Target produces high-quality genome assemblies by substantially reducing indel and mismatch errors, improving consensus quality, and increasing gene completeness. The tool also achieves these with lower computational costs compared to a current state-of-the-art utility.

## Implementation

### Algorithm Overview

The GoldPolish-Target workflow is implemented using SnakeMake (19) and consists of five stages (Figure 1). A draft sequence assembly and matching long-read sequences are required inputs. First, the long reads are mapped to the draft assembly with either ntLink (default) (18) or minimap2 (20), with the mappings output in PAF format (Pairwise mApping Format) (20). The PAF file format is a tab-delineated 12-column file that describes the mapping positions between two sets of sequences (21). Then, by default, GP-Target searches for soft-masked bases and excises them, providing a convenient means to polish raw bases left by the GoldRush pipeline.

**Figure 1.**
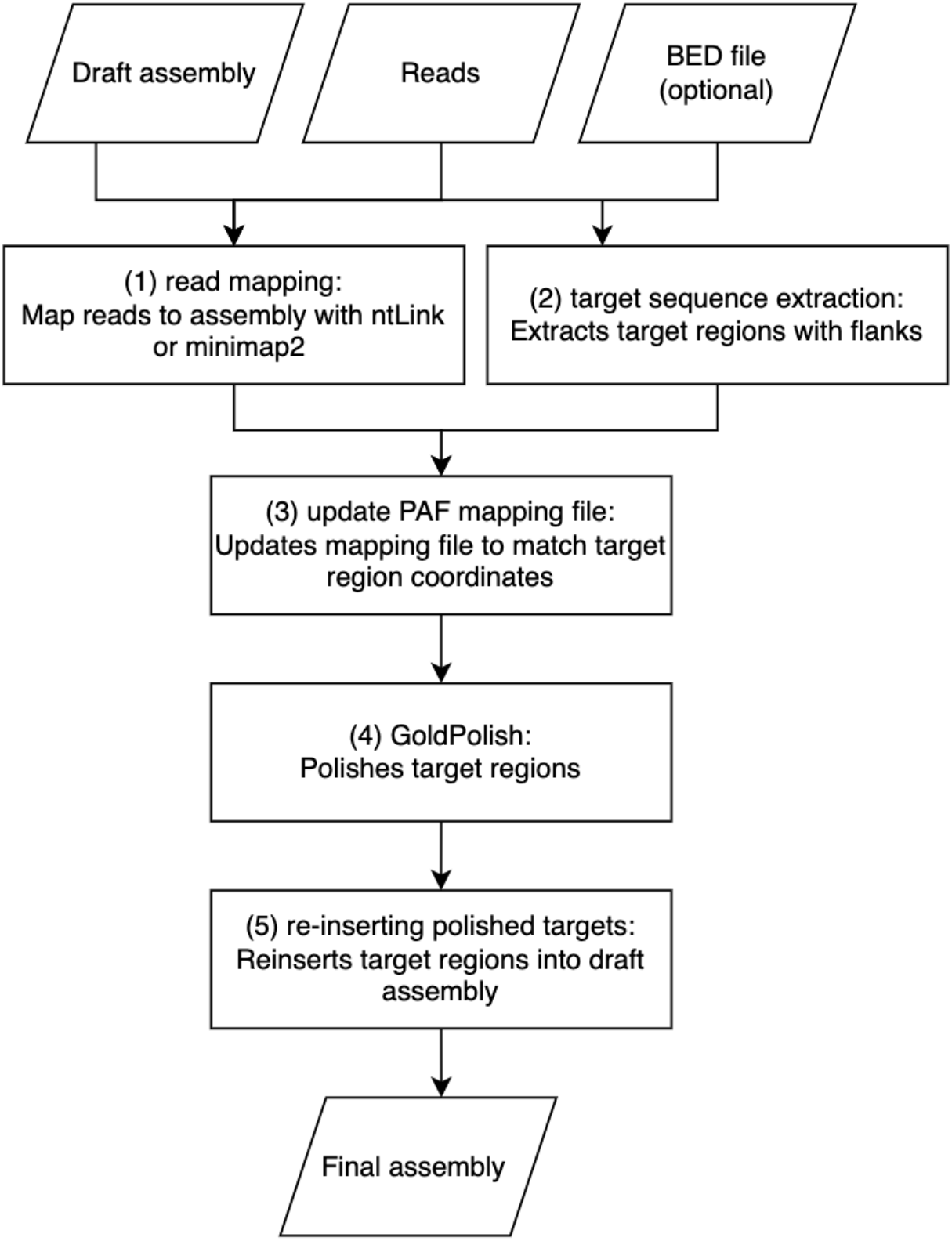
Schematic of GoldPolish-Target framework and steps. (1) Long-reads are first mapped to the draft assembly. (2) Then, by default, soft masked regions of the assembly are excised; alternatively, a BED file can be used to specify coordinates for the regions of interest to be excised. (3) The mapping file is modified to match the coordinate system of target regions and the resulting PAF file is passed to GoldPolish. (4) GoldPolish is run on the target regions to correct base errors. (5) The polished target sequences are re-inserted into the original assembly to yield the final polished assembly output.

Alternatively, users can submit a BED file with sequence coordinates for any number of target regions, specified using the chromosome, start position, and end position columns of the standard BED format (22). The sequences of the target regions are written to an intermediate FASTA file and the PAF mapping file generated by the mapping step is adjusted to match the naming and coordinate system of these target regions. Then, the updated PAF file and the intermediate FASTA files are passed to GoldPolish for polishing. Finally, the polished target regions are reinserted into the original draft assembly to yield the final polished assembly.

### Read mapping

By default, the long-read sequences are mapped to the draft assembly using the ‘pair’ function of ntLink (18). Users can optionally set the *k*-mer size (k_ntLink) and the window size (w_ntLink) for generating minimizers for the mapping; otherwise, these parameters will default to 88 bp and 1000 bp, respectively. Alternatively, minimap2 (20) with default parameters can be used for mapping.

### Target sequence extraction

The target regions are stretches of soft-masked bases in the draft assembly in the default mode. If a BED file is included as an input, GP-Target excises target sequences corresponding to the specified start and end positions. It extracts these target regions, plus flanking sequences 5’ and 3’ of the region (length; default 64 bp) and saves them to an intermediate FASTA file. When flanking sequences from multiple target regions overlap in the draft assembly, they are joined into one larger sequence in the resulting FASTA file. When added to the intermediate FASTA file, the names of the target regions are updated to include some key points of information. Each target region name begins with the name of the original contig from which it is extracted, followed by a numerical identifier denoting its relative position in the contig sequence (e.g., numbered 1, 2, …, n, where 1 denotes the first target region from the 5’ end of the sequence and ‘n’ represents the total count of target regions on that contig). Lastly, the start and end coordinates of these targets on the contig are appended to the FASTA header.

### Update PAF mapping file

The PAF file that is generated during the mapping step is updated to match the names of the target regions and their genomic coordinates; the start and end of the alignment block will be updated to map to the target region rather than the full, unpolished contig. Reads that do not align to any target region are discarded.

### GoldPolish code execution

Using the updated PAF file generated in step 3 (Figure 1) and the corresponding long-read sequences, GoldPolish is used to polish the intermediate FASTA file generated in step 2 (Figure 1). Using reads mapped to the draft assembly, GoldPolish generates Bloom filters, a succinct and probabilistic data structure (23), and populates them with words of length *k* (i.e. *k-*mers) from the long reads. These Bloom filters are used to correct mismatches and indels on each individual ‘goldtig’, a raw sequence in a 1X representation of the genome of interest, and close gaps between ‘goldtigs’ with uncorrected bases from a representative sequence read (11). A FASTA file with the polished target regions is generated as the final output of GoldPolish.

### Re-inserting polished sequence targets into the draft assembly

The polished sequence targets are re-inserted into the draft assembly using the coordinate information stored in their FASTA headers. This produces the final output FASTA file.

### Experimental data

ONT long reads from *Drosophila melanogaster*, sequenced using R10.4.1 pore chemistry, and the *Homo sapiens* NA24385 cell line, generated for the 1000 Genomes Project (24) sequenced using R9.4.1 pore chemistry, were obtained from the Sequence Read Archive (SRA) (Supplementary Table 1). For each set of reads, draft assemblies were generated using the standard GoldRush (v1.1.0) pipeline (11), with default parameters. To evaluate polishing with QUAST (25), we used a *D. melanogaster* reference genome retrieved from NCBI Reference Sequence Database (RefSeq) and the human genome reference GRCh38 from the Genome Reference Consortium (Supplementary Table 2). For evaluation with Merqury (26), we used Illumina HiSeq and NovaSeq reads sourced from the SRA for the *D. melanogaster* and *H. sapiens* assemblies, respectively (Supplementary Table 3).

### Comparison to the state-of-the-art

GoldPolish-Target (v1.0.0; default parameters) was compared to Medaka (v1.11.1; default parameters) (10). Identical BED files specifying the coordinates of the target regions in the draft assembly were passed to both polishing tools. Here, we targeted the soft-masked regions left by ntLink during the initial GoldRush assembly process; these are gaps that are filled with raw long reads that remain unpolished in the final draft assembly (18). The target regions were polished by Medaka in three separate steps: alignment of reads to the input assembly, running a consensus algorithm across assembly regions, and the aggregation of the subsequent results to create consensus sequences. Additionally, we ran Medaka as a global polishing tool as a comparator for GP-Target using the consensus feature. We assessed polishing results of both GP-Target and Medaka on the *D. melanogaster* and the *H. sapiens* assemblies.

### Machine specifications

All runs were benchmarked on a DELL server with 128 Intel(R) Xeon(R) CPU E7-8867 v3, 2.50 GHz with 2.6 TB RAM.

### Assessment of performance

The polished assemblies were assessed with QUAST (v5.0.2; --fast --large --scaffold-gap-max-size 100000 --min-identity 80 --split-scaffold) (25) and their corresponding reference genome (Supplementary Table 2). We used the number of mismatches and indels per 100 kbp to assess the base quality of the assemblies. Additionally, with the matching short-read paired-end sequences (Supplementary Table 3), Merqury was used (v1.3; default parameters) (26) to provide a reference-free, *k*-mer based assessment of the polished assemblies. We examined consensus quality value (QV) as a measure of the base quality changes introduced by polishing. Lastly, BUSCO (v5.3.2; default parameters) (27) was used to determine the number of protein-coding marker genes that were recovered by polishing to further assess the quality and completeness of the polished assemblies. To report average computational resource usage, time and peak random-access memory usage from triplicate runs were recorded.

## Results and Discussion

We assembled the genomes of *D. melanogaster* and *H. sapiens* using the GoldRush assembly pipeline. We then polished each of the draft assemblies with GP-Target (using ntLink and minimap2 mapping) and Medaka. The resulting polished assemblies were assessed using a series of quality metrics reported by QUAST (25), Merqury (26) and BUSCO (27), and their resource usage was also compared.

### Assessment of Base Error with QUAST

We compared the performance of GP-Target with Medaka in correcting insertion, deletion (indel), and mismatch errors in the draft assemblies using QUAST. Notably, even though the targeted regions in the *D. melanogaster* assembly constituted just 15% of the overall sequence by length of selected targets, GP-Target yielded substantial improvements in base accuracy. Specifically, it reduced indel errors by 49.2% (ntLink) and 43.5% (minimap2), and mismatch errors by 53.4% (ntLink) and 39.9% (minimap2) (Figure 2A). Polishing the same regions, Medaka produced 51.3% reductions in indels and 52.4% reductions in mismatches (Figure 2A). For the *H. sapiens* assembly, where only 5.2% of the assembly was targeted, GP-Target (ntLink) reduced indels by 16.4%, GP-Target (minimap2) by 16.5%, and Medaka by 18.1% (Figure 2B). Mismatch reductions were 14% (GP-Target ntLink), 13.3% (GP-Target minimap2), and 14% (Medaka) (Figure 2B). Here, both tools demonstrate comparable performance, showing a similar efficacy in fixing indels and base mismatch errors in the draft assemblies.

**Figure 2.**
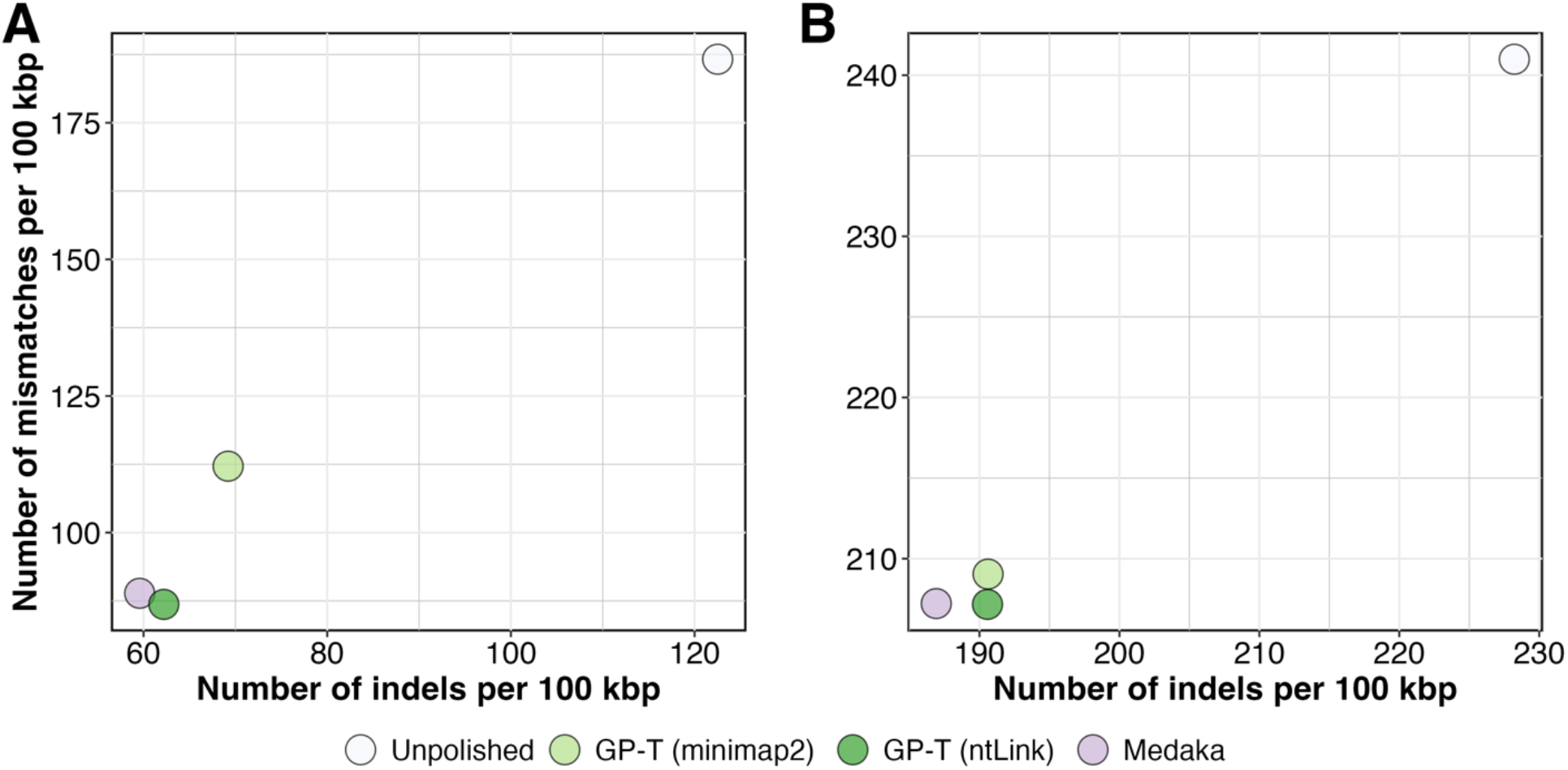
Indels and base mismatches before and after polishing GoldRush assemblies with GP-Target and Medaka. The draft assemblies were polished with GoldPolish-Target with minimap2 mapping, denoted as GP-Target (minimap2), GoldPolish-Target with ntLink mapping, denoted as GP-Target (ntLink), and Medaka with minimap2 mapping, denoted as Medaka. The numbers on the x- and y-axes, determined by QUAST, represent the number of indels per 100 kbp and the number of mismatches per 100 kbp before and after polishing the **A)** *D. melanogaster* and the **B)** *H. sapiens* draft GoldRush assembly.

### Assessment of Consensus Quality with Merqury

Next, we used Merqury, a reference-free, *k-*mer based assembly assessment tool, to assess consensus quality scores (QV) before and after the targeted polishing of each dataset. Alignment-based approaches, such as QUAST, can erroneously identify genuine variants as mismatches or indels, particularly in the absence of a high-quality or strain-specific reference genome to compare against (26). Merqury mitigates this reference bias by using *k-*mers derived from high-accuracy sequencing reads to assess assembly quality, rather than a reference genome (26). However, Merqury, while providing total error rates, does not differentiate indel and mismatch errors. As such, we used both QUAST and Merqury in conjunction to determine the impact of targeted polishing on assembly base quality. Merqury uses meryl, a k-mer counting tool, to estimate the frequency of consensus errors in the assembly (26). Through this process, Merqury estimates the consensus quality value, where a higher QV signifies a more accurate consensus—Q30 indicates 99.9% accuracy, Q40 denotes 99.99%, and so forth (26). Polishing the *D. melanogaster* assembly with GP-Target using ntLink and minimap2 mapping produced an 8.6% and 7.5% increase in assembly QV, respectively, compared to the 8.9% increase obtained with Medaka polishing (Figure 3A). This resulted in output assembly QV scores of 30.4, 30.1, and 30.5 for GP-Target (ntLink), GP-Target (minimap2), and Medaka, respectively (Figure 3A). Polishing the *H. sapiens* assembly, GP-Target demonstrated a 4.7% increase in assembly QV with both ntLink and minimap2 mapping, compared to a 5.1% increase with Medaka polishing (Figure 3B), corresponding to QV scores of 30.8 for both GP-Target polished assemblies and 30.9 for Medaka (Figure 3B). High QV scores represent an improvement over the appreciable error rates associated with ONT sequencing, with all polished assemblies achieving base accuracy exceeding 99.9%. QV increases are observed. These high QV scores are achieved after polishing just 15% and 5.2% of the *D. melanogaster* and *H. sapiens* genome assemblies based on the length of selected targets, respectively.

**Figure 3.**
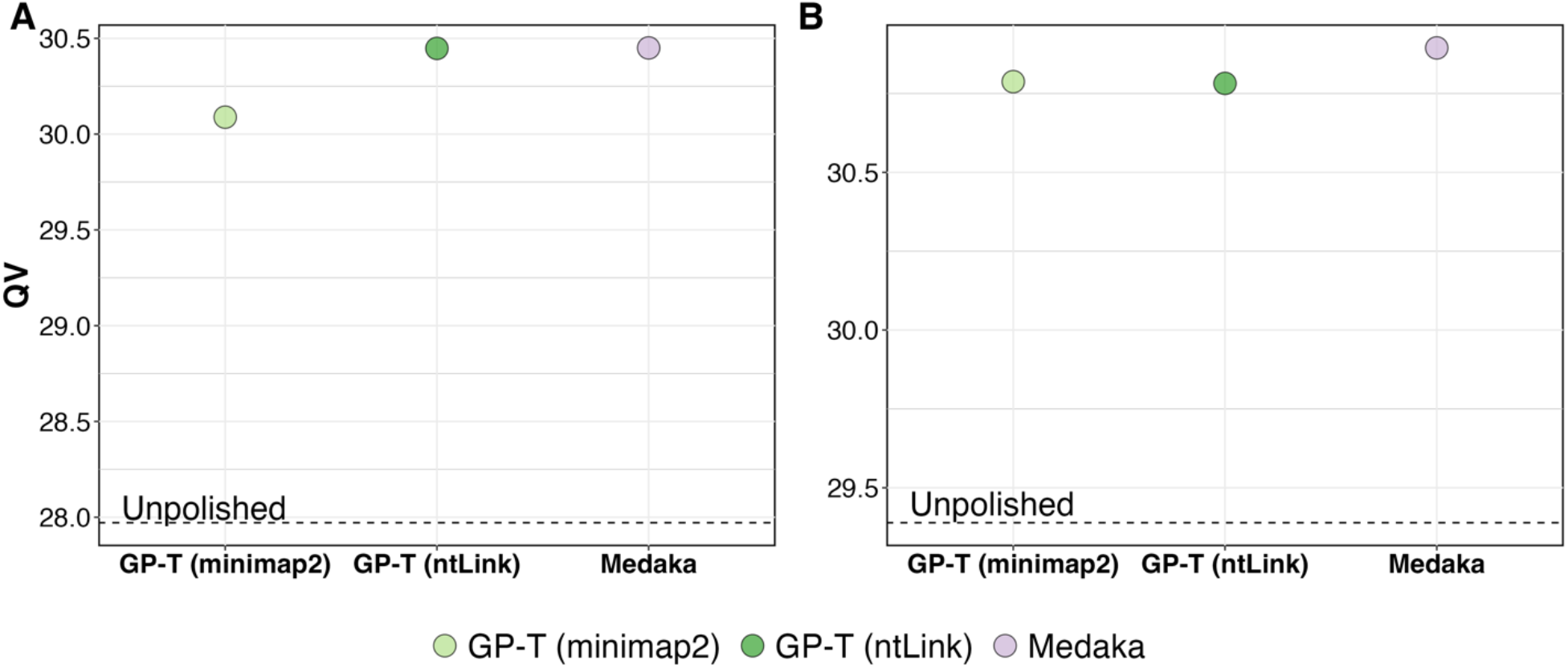
Merqury base error rate assessment of polished D. melanogaster and H. sapiens draft assemblies. The consensus quality (QV), as determined by Merqury, after polishing the **A)** *D. melanogaster* and **B)** *H. sapiens* draft GoldRush assemblies.

### Assessment of Gene Completeness with BUSCO

We used the Benchmarking Universal Single-Copy Orthologue (BUSCO) tool to assess the completeness of genome assemblies in the gene space (27). BUSCO detects the presence of evolutionarily-conserved single-copy genes within a lineage to assess the completeness of gene content within the genome (27). This evaluation method allows us to determine the capability of these polishing tools to fix mismatch and frameshift errors that could cause fragmented reconstruction in predicted gene ortholog products (27).

Assessing the *D. melanogaster* assembly, GP-Target’s polishing recovered seven complete BUSCOs (0.2% of all BUSCO groups surveyed) with both mapping options (Supplementary Table 4). Genes are classified as complete when their lengths are within two standard deviations of the BUSCO group mean length (27). In comparison, Medaka’s targeted polishing of the *D. melanogaster* draft assembly recovered nine complete BUSCOs (0.3% of all BUSCO groups surveyed) (Supplementary Table 4). For the *H. sapiens* dataset, the GP-Target polished assembly recovered 115 additional complete BUSCOs (0.8% of all BUSCO groups surveyed) with ntLink mapping and 67 (0.5% of all BUSCO groups surveyed) with minimap2 (Supplementary Table 5). Meanwhile, polishing with Medaka recovered 84 complete BUSCOs (0.6% of all BUSCO groups surveyed) (Supplementary Table 5). Despite targeting only a fraction of the *D. melanogaster* and *H. sapiens* assemblies, GP-Target’s correction of indel, mismatch, and frameshift errors aids in the recovery of additional of complete BUSCOs, ultimately producing more complete genome assemblies.

### Computational Resource Usage

While both GoldPolish-Target and Medaka produce polished genomes with comparable base quality and accuracy, GoldPolish-Target is more efficient in terms of total run time and peak random-access memory (RAM) usage regardless of the mapping algorithm used (ntLink or minimap2). When polishing smaller genomes, like *D. melanogaster*, GP-Target with ntLink mapping is the most efficient, running in 12.1 minutes with 1.4 GB peak RAM, on average— significantly faster and more resource-friendly than Medaka, which used an average of 3.4 hours and 33.8 GB of RAM (Figures 4A and 4B). Similarly, comparing GP-Target with minimap2 mapping to Medaka, GP-Target remains consistently more efficient, averaging 13.1 minutes and 3.8 GB of peak memory (Figures 4A and 4B).

**Figure 4.**
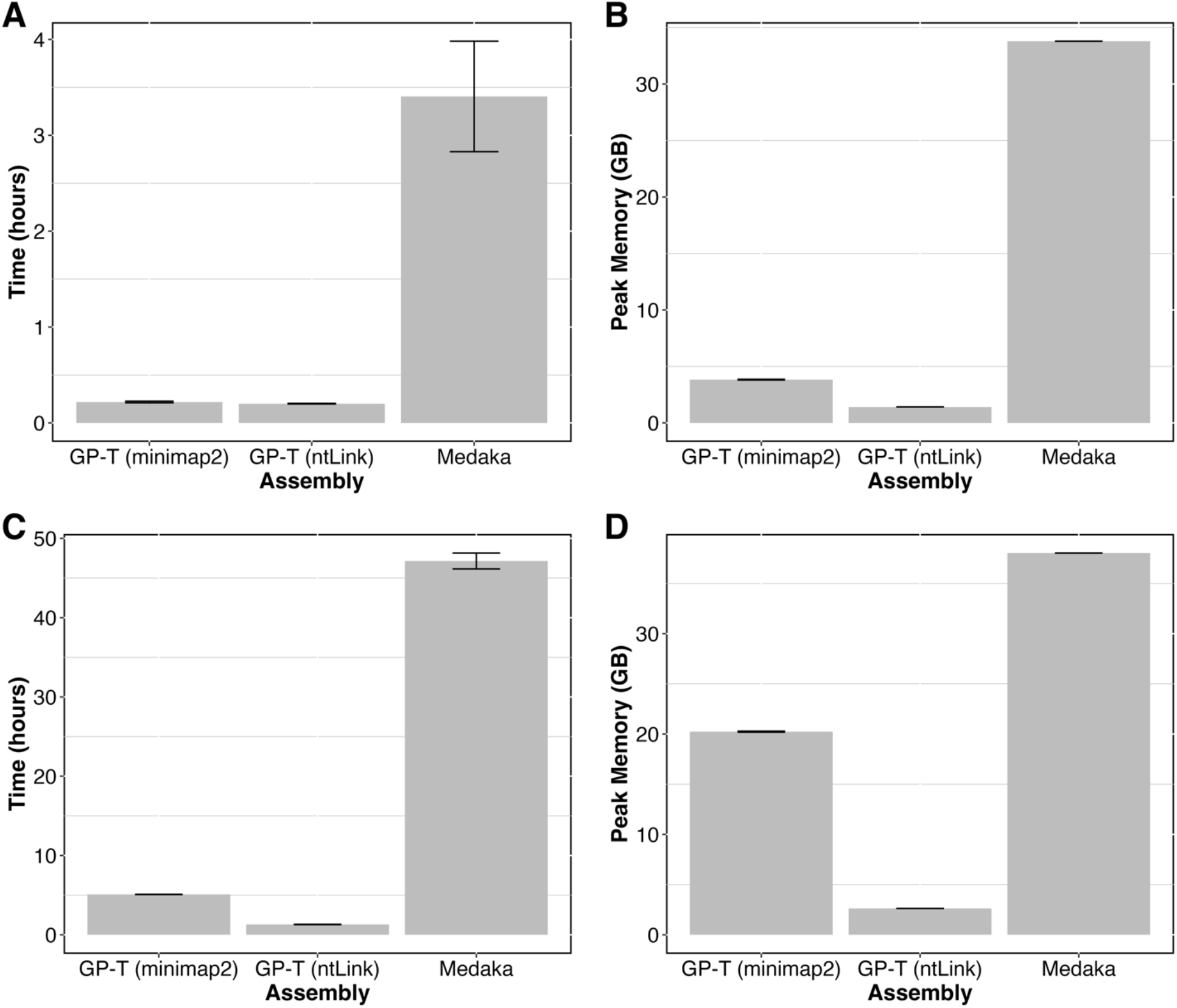
Compute resource usage of targeted polishing with GoldPolish-Target and Medaka. *D. melanogaster* draft assembly polishing **A)** wall-clock time (in hours) and **B)** peak memory (RAM, in gigabytes) and *H*. sapiens draft assembly polishing **C)** wall-clock time (in hours) and **D)** peak memory (RAM, in gigabytes) associated with GoldPolish-Target and Medaka. GoldPolish-Target with minimap2 alignment is denoted as GP-Target (minimap2) and GoldPolish-Target with ntLink mapping is denoted as GP-Target (ntLink).

GP-Target has a linear time complexity algorithm and scales favourably to large genomes, as shown by polishing the substantially larger *H. sapiens* assembly, which has a genome size 21.4-fold larger than that of *D. melanogaster*. With ntLink mapping, GP-Target achieves an average polishing time of 1.3 hours with a peak RAM usage of 2.6 GB with the *H. sapiens* dataset (Figures 4C and 4D). In contrast, Medaka requires 47.2 hours and 38.0 GB of peak RAM to polish the same target regions (Figures 4C and 4D). Comparing the two GP-Target mapping options, polishing with ntLink utilizes 12.9% of the peak RAM that minimap2 uses (Figure 4D).

When polishing the entire genome with Medaka, the base quality of both test assemblies was substantially reduced. Medaka, in its non-targeted mode, was able to decrease the number of mismatch errors by 79.7% and indel errors by 82.8% in the *D. melanogaster* assembly (Supplementary Figure 1). Moreover, it was able to reduce the number of mismatches and indels by 43.1% and 71.6%, respectively, for the *H. sapiens* assembly (Supplementary Figure 1). However, like its targeted mode, running Medaka as a global polisher required substantially more computational resources compared to GoldPolish-Target. Compared with GP-Target (ntLink), global polishing with Medaka spent 18.4X more time and 23.9X more RAM usage, on average, to polish the *D. melanogaster* assembly (Supplementary Figure 2). For the *H. sapiens* assembly, global polishing with Medaka used an average of 27.5X more time and 14.5X more memory (Supplementary Figure 2). While GP-Target does not improve the base quality of the two test genome assemblies as effectively as Medaka in its non-targeted mode, it offers a substantially more computationally lean polishing option, which has become increasingly vital to keep up with the onset of large-scale, high-throughput sequencing projects.

By specifying specific target regions, GP-Target offers a resource-efficient means of polishing genome assemblies. This allows users to target areas that are critical for analysis or highly erroneous regions, eliminating the need for polishing the entire assembly. Furthermore, by targeting specific regions on the draft assembly, GP-Target improves polishing accuracy by simplifying the Bloom filter-guided polishing step. Instead of generating one Bloom filter for each *k*-mer size per ‘goldtig’, GP-Target generates Bloom filters for each *k*-mer size per *target region*, only using reads mapped specifically to the target region, for error correction. When polishing draft assemblies with *D. melanogaster* and *H. sapiens* ONT long reads, GP-Target consistently reduces the number of indel and base mismatch errors, increases the quality value (QV), and recovers complete marker genes, as assessed by QUAST (25), Merqury (26), and BUSCO (27), respectively (Figures 2 and 3, Supplementary Tables 4 and 5). While Medaka shows similar or slightly better improvements in the same assembly metrics, GP-Target maintains a more modest computational resource usage for both test datasets (Figure 4).

In contrast, Medaka’s targeted functionality was intended to parallelize polishing the entire genome in separate regions rather than concentrating solely on targeted sections. Using Medaka for this approach required us to develop a BASH script to execute the three main steps separately—aligning reads, running the consensus algorithm, and combining the polished sequences together (10). GP-Target, on the other hand, requires minimal additional preparatory work.

GP-Target has been integrated as the last step in the GoldRush pipeline (v1.2.0) (11), automatically identifying and polishing gaps filled in with raw, unpolished reads by ntLink that are particularly prone to errors. This approach leads to substantial improvements in the base quality of the final output without having to polish the entire assembly. When using GP-Target to polish an assembly that is not soft-masked or for more controlled targeting of regions, users can instead generate a BED file with their desired polishing coordinates. Moreover, the consistently shorter run times associated with GP-Target polishing, facilitates quicker turnaround times, making it particularly advantageous for large-scale datasets and time-sensitive analyses.

## Conclusion

Here, we introduced GoldPolish-Target, a targeted long-read polishing pipeline designed to improve genome assembly base quality by targeting specific assembly regions. Our experiments, featuring fly and human datasets, demonstrate that GoldPolish-Target achieves polishing accuracy comparable to that of Medaka—a state-of-the-art polishing tool with a similar targeted functionality. However, GoldPolish-Target is consistently more computationally efficient, with shorter average run times and smaller memory footprints. Furthermore, GoldPolish-Target offers versatility, supporting mapping-based polishing through minimap2 or ntLink, while maintaining adaptability for potential integration with future tools. In summary, GoldPolish-Target emerges as a powerful, efficient, and flexible solution for targeted polishing genome assemblies using long reads.

