## Supplementary material for "GoldPolish-Target: Targeted long-read genome assembly polishing"

#### Supplementary Figures

**Supplementary Figure 1. Indels and base mismatches before and after polishing GoldRush assemblies with GP-Target and Medaka.** The draft assemblies were polished with GoldPolish-Target with minimap2 mapping, denoted as GP-Target (minimap2), GoldPolish-Target with ntLink mapping, denoted as GP-Target (ntLink), and Medaka in targeted mode, denoted as Medaka (targeted) and Medaka in whole genome polishing mode, denoted as Medaka (whole genome). The numbers on the x- and y-axes, determined by QUAST, represent the number of indels per 100 kbp and the number of mismatches per 100 kbp before and after polishing the **A)** *D. melanogaster* and the **B)** *H. sapiens* draft GoldRush assembly.

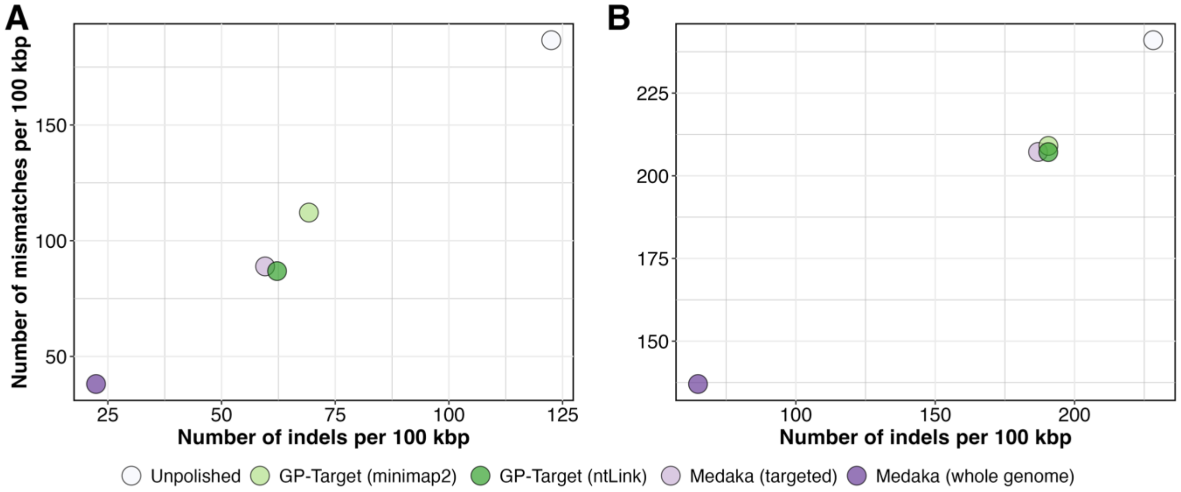

**Supplementary Figure 2. Compute resource usage of targeted polishing with GoldPolish-Target and Medaka**. *D. melanogaster* draft assembly polishing **A)** wall-clock time (in hours) and **B)** peak memory (RAM, in gigabytes) and *H.* sapiens draft assembly polishing **C)** wall-clock time (in hours) and **D)** peak memory (RAM, in gigabytes) associated with GoldPolish-Target and Medaka. GoldPolish-Target with minimap2 alignment is denoted as GP-Target (minimap2) and GoldPolish-Target with ntLink mapping is denoted as GP-Target (ntLink). Polishing only the target regions with Medaka is denoted as Medaka (targeted) and polishing the whole genome with Medaka is denoted as Medaka (whole genome).

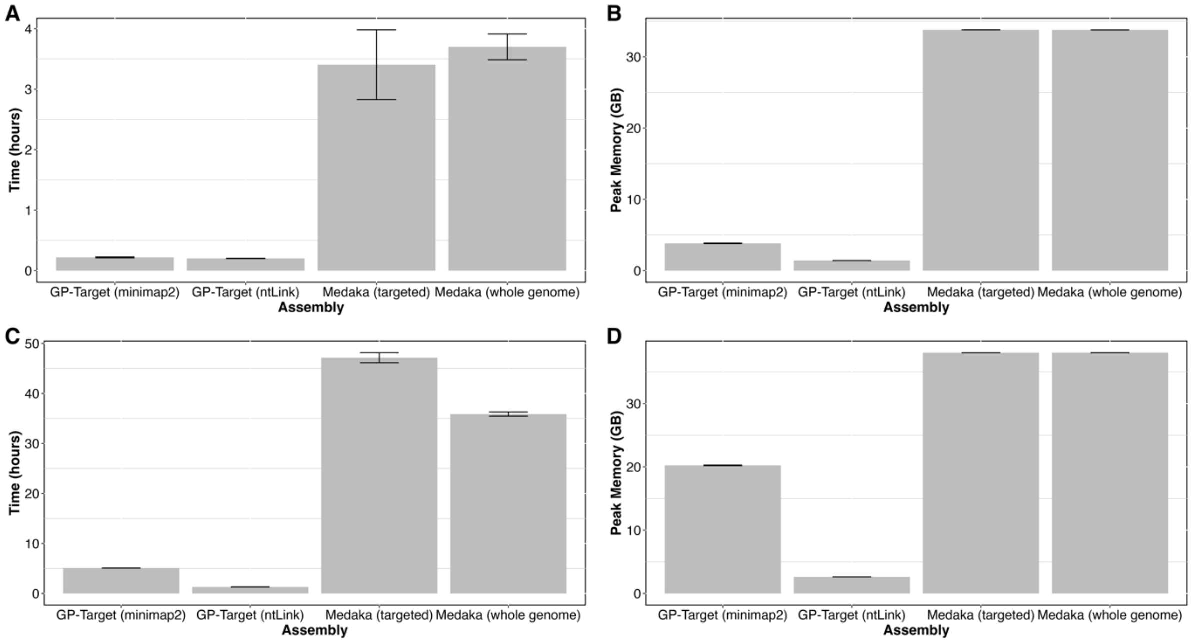

### Supplementary Tables

**Supplementary Table 1. ONT long-read sequencing reads used for genome assembly assessments**. The N50 length and associated base error rate for each data set were assessed with ABySS (1) and NanoSim (2), respectively.

| Species | Fold Coverage | N50 Length (bp) | Accession(s)/ Source | Basecaller | Flowcell | Estimated Error Rate (%) |
|---|---|---|---|---|---|---|
| *D. melanogaster* | 154 | 19,281 | SRR22822929 | Guppy v6 | R10.4.1 | 3 |
| *H. sapiens* | 67 | 30,348 | s3://ont-open-data/gm24385_2020.11/analysis/r9.4.1/20201026_1644_2-E5-H5_PAG07162_d7f262d5/guppy_v4.0.11_r9.4.1_hac_prom/align_unfiltered/chr1/guppy_v5.0.6_r9.4.1_sup_prom/ | Guppy v5 | R4.9.1 | 4 |

**Supplementary Table 2. Reference genomes used for QUAST assembly base quality evaluation**. Reference genomes for *D. melanogaster* and *H. sapiens* were sourced from the NCBI Reference Sequence Database and the Genome Reference Consortium (3), respectively.

| Species | Reference genome build | Accession |
|---|---|---|
| *D. melanogaster* | Release 6 | NC_004354 |
| *H. sapiens* | GRCh38 | GCA_000001405.15 |

**Supplementary Table 3. Short read datasets used for Merqury assembly base quality evaluation**.

| Species | SRA Accession |
| --- | --- |
| *D. melanogaster* | SRR11460799 |
| *H. sapiens* | SRR11321732 |

**Supplementary Table 4. BUSCO statistics for the *D. melanogaster* assembly before and after polishing.** BUSCO was run with diptera_odb10 lineage and 3,285 BUSCO groups were searched (4).

| Assembly | Complete BUSCOs (C) | Complete and single-copy BUSCOs (S) | Complete and duplicated BUSCOs (D) | Fragmented BUSCOs (F) | Missing BUSCOs (M) |
| --- | --- | --- | --- | --- | --- |
| Unpolished | 3,219 (98.0%) | 3,164 | 55 | 31 | 35 |
| Medaka | 3,228 (98.3%) | 3,106 | 122 | 27 | 30 |
| GP-T (minimap2) | 3,226 (98.2%) | 3,099 | 127 | 29 | 30 |
| GP-T (ntLink) | 3226 (98.2%) | 3,101 | 125 | 28 | 31 |

**Supplementary Table 5. BUSCO statistics for the *H. sapiens* assembly before and after polishing.** BUSCO was run with primates_odb10 lineage and 13,780 BUSCO groups were searched (4).

| Tool | Complete BUSCOs (C) | Complete and single-copy BUSCOs (S) | Complete and duplicated BUSCOs (D) | Fragmented BUSCOs (F) | Missing BUSCOs (M) |
|---|---|---|---|---|---|
| Unpolished | 12,238 (88.8%) | 12,025 | 213 | 501 | 1041 |
| Medaka | 12,322 (89.4%) | 12,028 | 294 | 502 | 956 |
| GP-T (minimap2) | 12,305 (89.3%) | 11,988 | 317 | 507 | 968 |
| GP-T (ntLink) | 12,353 (89.6%) | 12,042 | 311 | 469 | 958 |
